# Field-Sensitive Axon Initial Segment Polarization by Ionic Direct Current Dissociates Gain Modulation from Temporal Structure in Cortical Circuits

**DOI:** 10.1101/2025.06.19.660597

**Authors:** Runming Wang, Gene Fridman

## Abstract

Conventional implantable neuromodulation relies on brief electrical or optical pulses that directly evoke action potentials in targeted neurons. While powerful, such approaches primarily impose activity rather than modulate how circuits process ongoing signals. Here, we introduce ionic direct current (iDC) delivered through non-penetrating, electrolyte-filled microcatheters on the cortical surface as an alternative strategy for neuromodulation that adjusts cortical gain without directly driving spikes. In rat primary somatosensory cortex, laminar recordings showed that cathodic iDC attenuated and anodic iDC amplified spontaneous and sensory-evoked activity while preserving the temporal structure of population responses. Computational modeling implicated polarization at the axon initial segment as the mechanism underlying bidirectional gain control. In awake animals, iDC modulation altered tactile sensitivity, demonstrating behavioral relevance. These findings establish iDC as a rapidly reversible, spatially precise method for modulating cortical processing, offering a fundamentally different mode of neuromodulation that bridges circuit physiology and behavior without overriding native neural dynamics.

**TEASER:** Ionic direct current modulates cortical gain via subthreshold polarization, preserving native neural dynamics.

## INTRODUCTION

Graded approaches to testing causal involvement can complement binary measures of necessity or sufficiency in studies of how neural circuits contribute to behavior. We introduce ionic direct current (iDC) neuromodulation (*1–4*) as a method to bidirectionally tune cortical gain with spatial and temporal precision. We test the hypothesis that iDC neuromodulation can scale neural population responses in vivo without altering their temporal structure, and we examine behavioral relevance by altering tactile sensitivity in awake animals.

Traditional circuit-dissection methods establish necessity through lesions or silencing, and sufficiency through stimulation (**Fig. 1A**). These approaches, implemented with techniques such as electrical microstimulation(*5*, *6*) or optogenetics(*7*, *8*), are powerful, but their pulsatile control can impose time-locked synchrony that overwrites native dynamics and has limited ability to quantify the *degree* of involvement. Many natural cortical computations evolve in high-dimensional, context-dependent population activity in which information is carried both by firing rates and by the relative timing and correlation structure across neurons(*9–12*). In such regimes, phase-locked stimulation may be a poor match to the underlying code. Drug-based gain modulation (receptor agonists or antagonists) can apply a tonic bias to probe the degree of network involvement by shifting excitability, but it lacks spatial and temporal precision (**Fig. 1B**).

**Figure 1.**
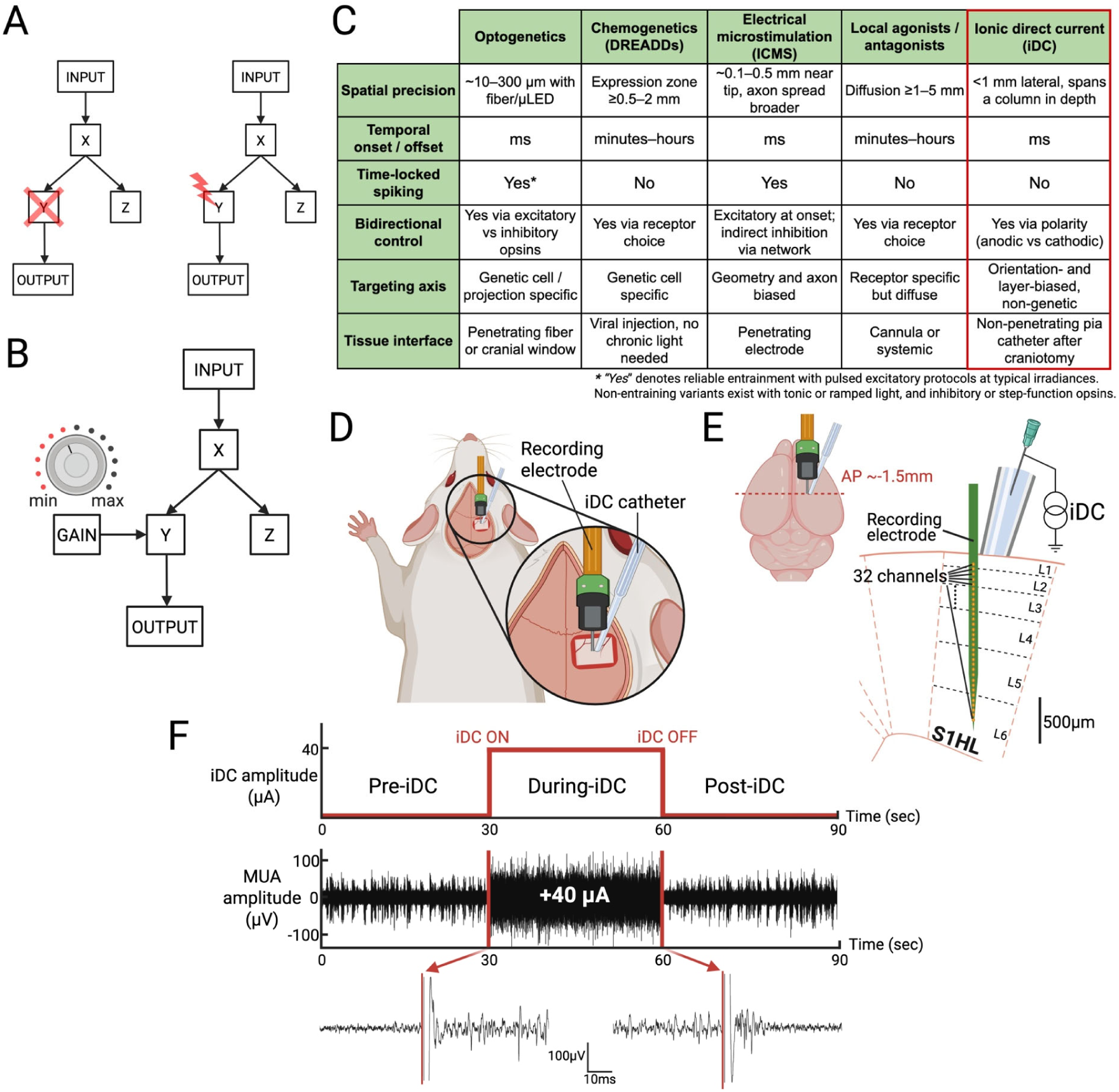
**A**) Simplified hypothetical set of brain neural modules X, Y, Z that are affected by experimental input. Experimental Input causes X to respond, which causes Y and Z, but only Y causes the OUTCOME. *left:* Ablating or temporarily inhibiting a module examines its necessity toward the behavioral outcome. *right:* Exciting neural activity at the output of a module <u>may</u> examine the causal effect on the outcome. **B)** Modulating the gain of a module examines the degree to which the module causally contributes to the outcome. **C)** Comparison of neuromodulation modalities on six axes: spatial footprint, onset and offset, entrainment risk, bidirectionality, targeting axis, and tissue interface. Values are typical and parameter dependent. iDC refers to separated-interface electrolyte direct current at the pia via a non-penetrating microcatheter. **D)** iDC stimulation catheter and recording electrode setup in acute rat S1HL experiments. **E)** Scaled cartoon of the 32-channel recording electrode placed within the six layers of the S1HL cortex. **F)** *top:* iDC stimulation timeline when recording spontaneous neural activity. *middle:* Temporally aligned representative MUA recording with +40µA anodic iDC stimulation from one electrode channel. *bottom*: Magnified MUA signal snippets during iDC onset and offset.

Electric fields arise naturally from neural activity and can in turn influence it (*13–15*). Artificially created electric fields therefore could offer an alternative to evoking spikes directly with all-or-nothing approaches. It has been shown in vitro that weak fields subtly polarize neuronal membranes without driving synchronized spikes, shifting how excitatory and inhibitory inputs are integrated(*16–18*). However, the approach of delivering sustained current to stimulate neurons in vivo has been impractical since metal electrodes implanted in the body will generate electrochemical reactions for pulses longer than several hundred microseconds(*19*, *20*). The recent development of the separated nerve interface and the Freeform Stimulator (FS) implant removes these constraints(*21*, *22*) by isolating metal components from tissue and allows arbitrary electric fields (including iDC) to be delivered indefinitely at the cortical surface. This architecture provides a practical substrate for chronic implementations of iDC-based cortical implants.

A comparative view clarifies where iDC fits among existing tools (**Fig. 1C**). iDC delivered through electrolyte microcatheters is hypothesized to provide millimeter-scale precision with millisecond onset and offset, preserve native time coding, and support bidirectional and rapidly reversible control via polarity. Because current is delivered locally at the pia through a small outlet, this localized gain control limits recruitment of distant regions, improves causal attribution at the mesoscale, and enables graded tests of necessity within a targeted cortical unit. Although the catheter does not penetrate, field lines couple through the cortical depth, so a single surface source can bias an entire column.

Here we chose a well-understood cortical circuit to investigate the neuromodulatory effects of iDC delivered through electrolyte microcatheters in rat S1HL. We show polarity– and layer-dependent scaling of spontaneous and sensory-evoked activity while preserving intrinsic spatiotemporal patterns. A biophysical model links these effects to field-sensitive dendritic summation near the axon initial segment with layer-specific consequences. Lastly, we demonstrate behavioral relevance by shifting tactile sensitivity thresholds in awake animals.

## RESULTS

### Modulation of Spontaneous Neural Activity

To investigate how ionic direct current (iDC) modulates spontaneous neural activity, we conducted a series of in vivo experiments in which a 32-channel single-shank microelectrode array was vertically inserted 1.65 mm into the S1HL cortex of urethane-anesthetized rats, spanning all cortical layers over a 1.55 mm contact array. A 250 µm-ID iDC microcatheter filled with electrolytic gel was then positioned on top of the pia mater, with its center located at a fixed horizontal distance (0.2 mm) from the recording electrode (**Fig.1DE**).

Spontaneous multi-unit activity (MUA) was recorded in ∼90-second sessions at different iDC amplitudes, each segmented into 30 s pre-iDC, 30 s during-iDC, and 30 s post-iDC epochs (**Fig. 1F, top**). A representative 90s multiunit activity (MUA) recording from a single channel with +40 µA iDC stimulation is shown (**Fig. 1F, middle, bottom**). The magnified snippets illustrate that a clear transition in spontaneous neural activity occurs immediately after the onset and offset of iDC stimulation.

To robustly quantify the spontaneous activity across channels and amplitudes, the MUA signals were processed using the Entire Spiking Activity (ESA) method (*23–25*). This processing method avoids threshold-based spike detection and yields a continuous measure of neural activity that is less sensitive to random changes in signal-to-noise ratio while retaining the spike contribution from smaller neurons (**Fig. 2A**) (*23*, *24*). Because the 32 channels spanned all cortical layers, we constructed depth-resolved ESA heat maps at each iDC amplitude to examine how iDC affected spontaneous activity across time and cortical depth (**Fig. 2B**). The heat maps show that before iDC stimulation is delivered (baseline, 0µA iDC), the most prominent activity happens in deeper layers, revealing robust slow wave oscillatory dynamics (∼1 Hz) characteristic of the urethane-anesthetized state (*26*, *27*). At lower iDC amplitudes (–30 to +30 µA), the slow wave oscillation remains intact, with only amplitude modulations in a graded fashion. In contrast, at higher iDC amplitudes (beyond ±30 µA), cathodic iDC produces near-complete suppression of activity in deeper layers, while anodic iDC elicits a persistent, high-frequency firing pattern. These findings suggest an approximately linear regime of graded modulation at moderate amplitudes and a nonlinear transition at higher amplitudes where iDC begins to suppress or drive activity beyond endogenous oscillatory patterns.

**Figure 2.**
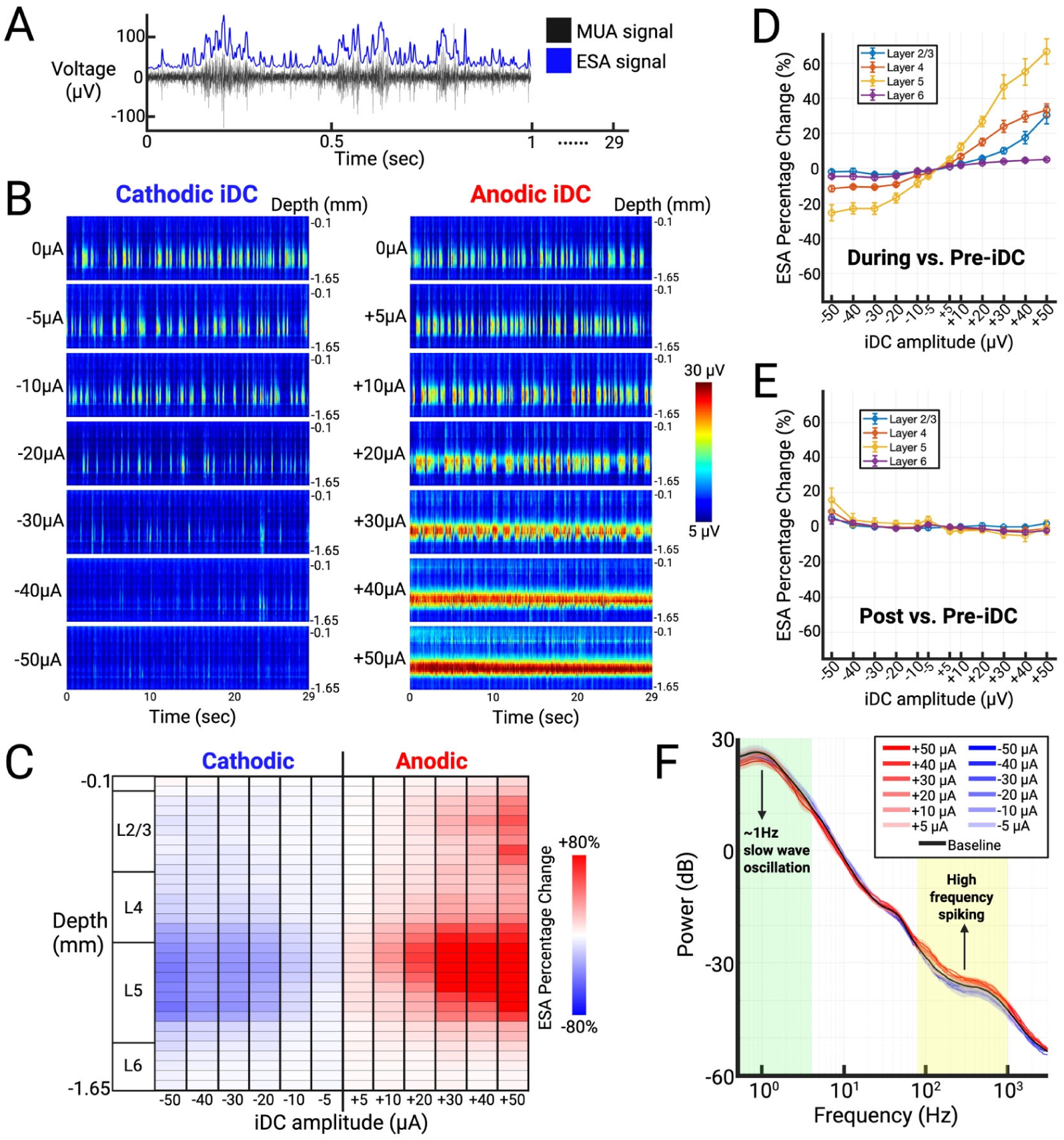
**A**) Representative MUA snippet (black) overlaid with processed Entire Spiking Activity (ESA) signal (blue). **B)** Time-resolved ESA heat maps across iDC amplitudes (±5 to ±50 µA) from one representative experiment. **C)** Relative percent change in average ESA (during– vs. pre-iDC) across recording channels over a range of iDC amplitudes from the same experiment as B. **D)** Cumulative ESA percent change (during– vs. pre-iDC) grouped by layers L2/3, L4, L5, and L6 (n = 7 animals, mean ± SEM). **E)** Cumulative ESA percent change (post– vs. pre-iDC) grouped by layers L2/3, L4, L5, and L6 (n = 7 animals, mean ± SEM), showing rapid reversibility after iDC offset. **F)** PCA-derived column power spectral density (PSD) across iDC amplitudes and pre-iDC baseline (n = 3 animals, mean ± SEM).

To summarize the relative effects of iDC across layers, we computed the percent change in averaged ESA between pre-iDC and during-iDC epochs for each channel and grouped channels by cortical layer (**Fig. 2C**). The most pronounced modulation occurred in layer 5 (L5), with the magnitude of the effect increasing with iDC amplitude. For anodic stimulation specifically, iDC also induced a gradual increase in layer 2/3 (L2/3) activity with higher amplitudes.

To determine whether this laminar pattern generalized across animals, we compiled results from all seven rats that underwent the spontaneous iDC experiments. In **Figure 2D**, the ESA percent change (during– vs. pre-iDC) values were averaged across channels within each layer, with the standard error of the mean (SEM) across multiple animals (n=7). The line plot shows that L5 is indeed the most affected layer across experiments, followed by layers 2/3 and 4 (L4), whereas layer 6 (L6) showed only minimal changes across the same range of iDC amplitudes. Across layers, response magnitude scaled approximately linearly with iDC amplitude between −30 and +30 µA, delineating the dynamic range over which iDC modulates network excitability without fully suppressing or saturating activity.

To assess reversibility, we computed post– vs. pre-iDC ESA percent change immediately after stimulation offset (**Fig. 2E**). Across layers and amplitudes, values clustered near zero, indicating rapid return to baseline once iDC was removed. A small residual offset was apparent at the largest cathodic setting (−50 µA), most evident in L5 and accompanied by larger SEM, while all other amplitudes remained within a few percent of zero. These data show that the modulation produced by iDC resolves promptly once the current ceases across the stimulation range tested.

To confirm that iDC modulates local neuronal gain without fundamentally altering the global oscillatory state, we performed a complementary frequency-domain analysis by comparing power spectral density (PSD) curves generated from unfiltered raw signals, rather than ESA, to eliminate preprocessing effects. In **Figure 2F**, the PSD curves are shown for a range of iDC amplitudes relative to the baseline recording, with the shaded region indicating SEM across multiple animals (n=3). Due to reference-wire artifacts affecting raw signal quality in four animals, PSD analyses were restricted to the three datasets where this issue was absent. These artifacts did not affect ESA metrics (see **Supplemental Fig. 1**), supporting the reliability of the broader findings. Across all tested amplitudes from −50 to +50 µA, the ∼1 Hz slow-wave peak characteristic of the urethane-anesthetized state was preserved relative to the pre-iDC baseline, indicating that the global oscillatory dynamics remained intact. Consistent with our ESA findings, the higher-frequency components (∼80–1000 Hz) of the PSD curves, which more directly reflect local spiking activity (*28*), exhibited clear polarity– and amplitude-dependent spectral power changes at high amplitudes.

### Computational Model and Experimental Assessment of iDC Modulation

Motivated by the laminar, polarity-dependent gain modulation observed in spontaneous ESA recordings, we next asked whether these effects could be explained by a field-induced potential bias toward inhibition or excitation at the axon initial segment (AIS) summation point of the pyramidal neurons. To address this, we built a computational model of field-sensitive AIS polarization in rat S1HL that simulates the extracellular electric field produced by iDC and the resulting membrane polarization along the somatodendritic axis of different neuron classes. The model first establishes a two-dimensional grid representing a cortical cross-section and calculates the extracellular potential (Vₑ) generated by a 250 µm-diameter isopotential disk (mimicking the iDC catheter) placed on the pia mater. This yields an estimate of the extracellular voltage Vₑ (relative to distant reference) along the cortex and in-depth along the cortical layers (**Fig.3A**).

**Figure 3.**
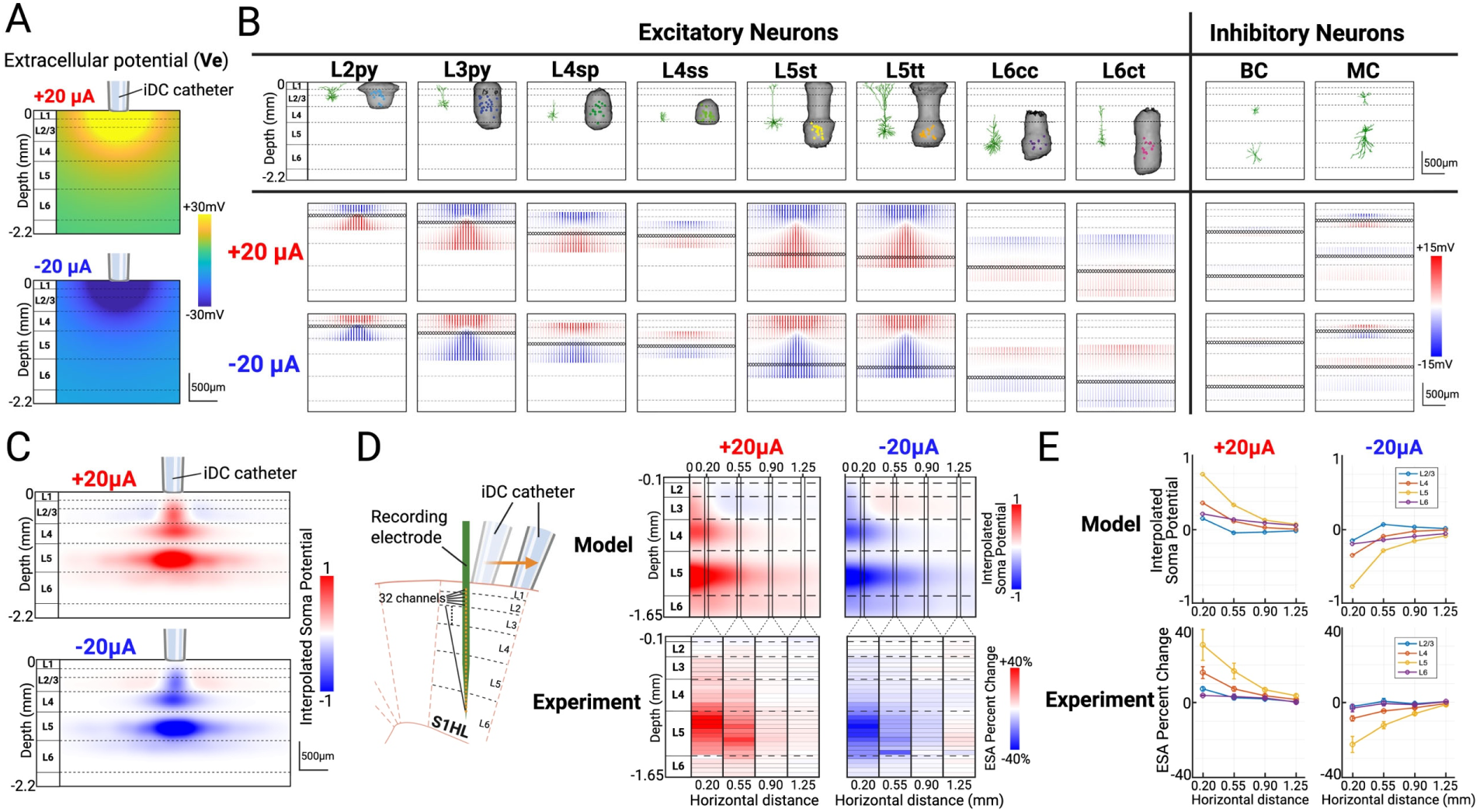
**A**) Extracellular potential within a two-dimensional cortical cross-section generated by the iDC microcatheter placed at the top of the pia mater. **B) First row:** representative morphologies and dendrite 3D volumes of the 8 most common types of excitatory neurons (L2py: L2 pyramidal cell; L3py: L3 pyramidal cell; L4sp: L4 star pyramidal cell; L4ss: L4 spiny stellate cell; L5st: L5 slender-tufted pyramidal cell; L5tt: L5 thick-tufted pyramidal cell; L6cc: L6 corticocortical pyramidal cell; L6ct: L6 corticothalamic pyramidal cell. Modified from Radnikow and Feldmeyer, 2018, Narayanan et al., 2017, and Oberlaender et al., 2012) and the 2 most common types of inhibitory interneurons (L4 and L6 basket cells (BC); L2/3 and L5 Martinotti cells (MC). Modified from Feldmeyer et al., 2018) within the rat somatosensory cortex. **Second and third row:** Each major neuron type is being modelled as vertical rods placed within the cortex cross-section, and their soma locations are shown as black circles. The color denotes the mirror estimate of the membrane potential change along the vertical length of each neuron in response to the extracellular electric field induced by anodic +20µA and cathodic –20µA iDC stimulation, respectively. **C)** The weighted sum of all membrane potential changes at excitatory neuron somas in proportion to their relative densities. **D)** Predicted iDC-induced somatic potential changes based on computational model versus representative experimental result by horizontally moving the iDC catheter to specific distances from the recording electrode (0.20, 0.55, 0.90, and 1.25mm). **E)** Cumulative model and ESA percent change versus horizontal distance plot from multiple animals (n=5, mean ± SEM).

In our model, eight principal excitatory neuron types (layer 2/3 pyramidal (L2py, L3py), layer 4 spiny stellate (L4sp), layer 4 star pyramidal (L4ss), layer 5 slender-tufted (L5st), layer 5 thick-tufted (L5tt), layer 6 corticocortical (L6cc), and layer 6 corticothalamic (L6ct)) and two common inhibitory neuron types (basket cells (BC) and Martinotti cells (MC)) are parameterized by their top and bottom dendritic depths as well as their soma depths according to their biological morphologies (**Fig.3B, first row**) (*29–32*). Neurons are modeled as simplified vertical one-dimensional ‘rods’ whose lengths span the extent of their dendritic arbors along the somatodendritic axis, and each rod includes a defined soma (indicated by a black marker) that represents the location of the cell body. For each neuron, we then compute a “mirror estimate” of membrane potential change along the rod when it is exposed to the electric field. This is defined as the difference between the mean extracellular voltage 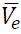 along the vertical extent of the cell and the local *V*_*e*_ at each position (*2*, *33*, *34*). The lower two rows of **Figure 3B** show each neuron as a single rod with the color indicating the membrane potential change along the length of the neuron. These are shown at incremental lateral positions across the cortical plane to illustrate how the neuron membrane voltages change across the horizontal distance under ±20 µA stimulation. In our analysis, we treat the modeled soma position as a proxy for the axon initial segment (AIS), which in these cell types lies within tens of microns of the soma. Because the extracellular field varies smoothly on this spatial scale, the predicted polarization at the soma marker provides a reasonable approximation to AIS polarization.

According to the model, L5 pyramidal cells with long somatodendritic axes (e.g., L5st, L5tt) display the most pronounced somatic depolarization under anodic (+20 µA) iDC currents. These neurons are large projection cells with long apical dendrites extending toward the cortical surface and somata positioned near the base of the dendritic volume, so that their somatodendritic axis is approximately aligned with the radial field generated by the pia-mounted iDC source. Under anodic current, superficial apical dendrites lie in a relatively hyperpolarized region along the neuron, whereas deeper basal dendrites and the soma reside in a depolarized region, yielding a large mirror-estimate gradient along the cell with a positive extremum at the soma proxy. Because action potentials are initiated at the AIS, which lies close to the soma and contains a high density of voltage-gated sodium channels, this predicted depolarization is expected to lower spike threshold and increase L5 firing probability. Cathodic (−20 µA) iDC inverts the field pattern, placing the soma/AIS proxy in a hyperpolarized region, raising threshold, and suppressing activity in these neurons. By contrast, L2/3 pyramidal neurons (e.g., L2py, L3py), though vertically oriented, have shorter apical arbors whose medial soma positions lie near the biphasic zero-crossing of the field, resulting in minimal net polarization effects. L4 neurons (e.g., L4sp, L4ss) yield similarly attenuated effects. L6 pyramidal cells (e.g., L6cc, L6ct), despite long apical extensions and somas near their basal dendrites, experience reduced field strength because the end of their apical dendrites does not reach as superficially as L5 pyramidal cells.

The two most common types of inhibitory interneurons (basket cell and Martinotti cell) were also modeled accordingly (**Fig.3B, right**) (*31*). Many cortical inhibitory interneurons, especially basket cells, exhibit compact, roughly symmetric dendritic arbors. Moreover, because inhibitory neurons make up a smaller fraction of the neuronal population (15-20%) and their synaptic outputs are more diffusely targeted, we expect the direct effects of iDC stimulation on these interneurons to be minimal (i.e. close to net zero) compared to the robust modulation seen in deep pyramidal cells (*35*). Thus, the model predicts that the primary direct effect of iDC at the cellular level is a polarity-dependent shift in AIS membrane potential in deep pyramidal neurons, with only modest net polarization in more superficial pyramidal cells and in most interneurons.

To mimic the spatial variability found in the cortex, rods representing each cell type are randomly scattered across the horizontal axis without overlapping, with the number of rods per type scaled according to their reported densities (*30*). The model then extracts the membrane potential change at each soma and applies a Gaussian-weighted sum interpolation over a finer grid. This produces a continuous two-dimensional heatmap that reflects the cumulative effect of iDC stimulation on the local population of neuronal activity. Since the AP threshold is determined primarily at the axon initial segment (AIS) due to the high density of voltage-gated sodium channels, and since the recording electrode primarily picks up somatic spiking activity, this interpolated potential change heatmap serves as an approximation of the local “gain kernel” that iDC stimulation would apply onto the electrophysiology signals of the cortical cross-section (**Fig. 3C**). The resulting figure indicates that the main depth of depolarization and hyperpolarization is predicted to be around L5, with a weaker modulation effect seen in L2/3 and L4, which progressively decays as horizontal distance increases. Interestingly, the model also predicted a faint, reversed polarity band in L2/3 at distances >0.2 mm from the iDC catheter.

Building on the model’s predicted gain kernel, we next tested the spatial resolution of iDC modulation in vivo using the same acute recording setup described in **Fig.1C**. To probe lateral field spread without moving the recording electrode, which is experimentally prohibitive, we shifted the iDC catheter horizontally along the pia to offsets of 0.20, 0.55, 0.90, and 1.25 mm from the recording electrode (**Fig. 3D, left**). Spontaneous MUA was recorded in 90-second runs, each divided into 30 s pre-iDC, during-iDC, and post-iDC epochs. The relative percent change between the average ESA for pre-iDC and during-iDC epoch was computed for each channel from a representative experiment, which forms a table with rows representing individual channels and columns corresponding to different lateral distances of the iDC microcatheter (**Fig. 3D, right**). Anodic modulation (+20 µA) produced maximal ESA increases in L5 that diminished with distance, effectively vanishing beyond ∼1 mm, which confirms the millimeter-scale spatial resolution of iDC modulation. Cathodic stimulation (–20µA) yielded complementary decreases. These patterns in spatial decay observed in experiments closely align with our model’s predictions, supporting the idea that the millimeter-scale gain field seen in the data arises from focal AIS polarization in deep pyramidal neurons under iDC. Interestingly, we note a faint reversed-polarity modulation in some L2/3 channels at larger lateral offsets, consistent with a predicted sign inversion near the field zero-crossing. A cumulative ESA percent change versus horizontal distance plot from multiple animals (n=5, mean ± SEM) suggests that this pattern is consistent across repetitions (**Fig. 3E**).

### Modulation of Foot Stimulation-Evoked Responses

To assess how iDC modulates the gain of evoked responses in S1HL, we extended the experiment described in **Figure 1C** to deliver controlled foot shock stimulation while delivering iDC on top of the pia mater (**Fig. 4A**). We first confirmed recording electrode placement by gently brushing the contralateral hindfoot, which elicited robust spiking responses across our 32-channel array. Foot stimulation was delivered through a surface electrocardiogram electrode using 200 µs cathodic pulses at 2-second intervals (10 repetitions per epoch) in a 90-second recording structure, comprising 30 s pre-iDC, 30 s during-iDC, and 30 s post-iDC epochs (**Fig. 4B**). For control, the same stimulation protocol was applied to the ipsilateral foot, confirming that the observed responses were specific to contralateral stimulation.

**Figure 4.**
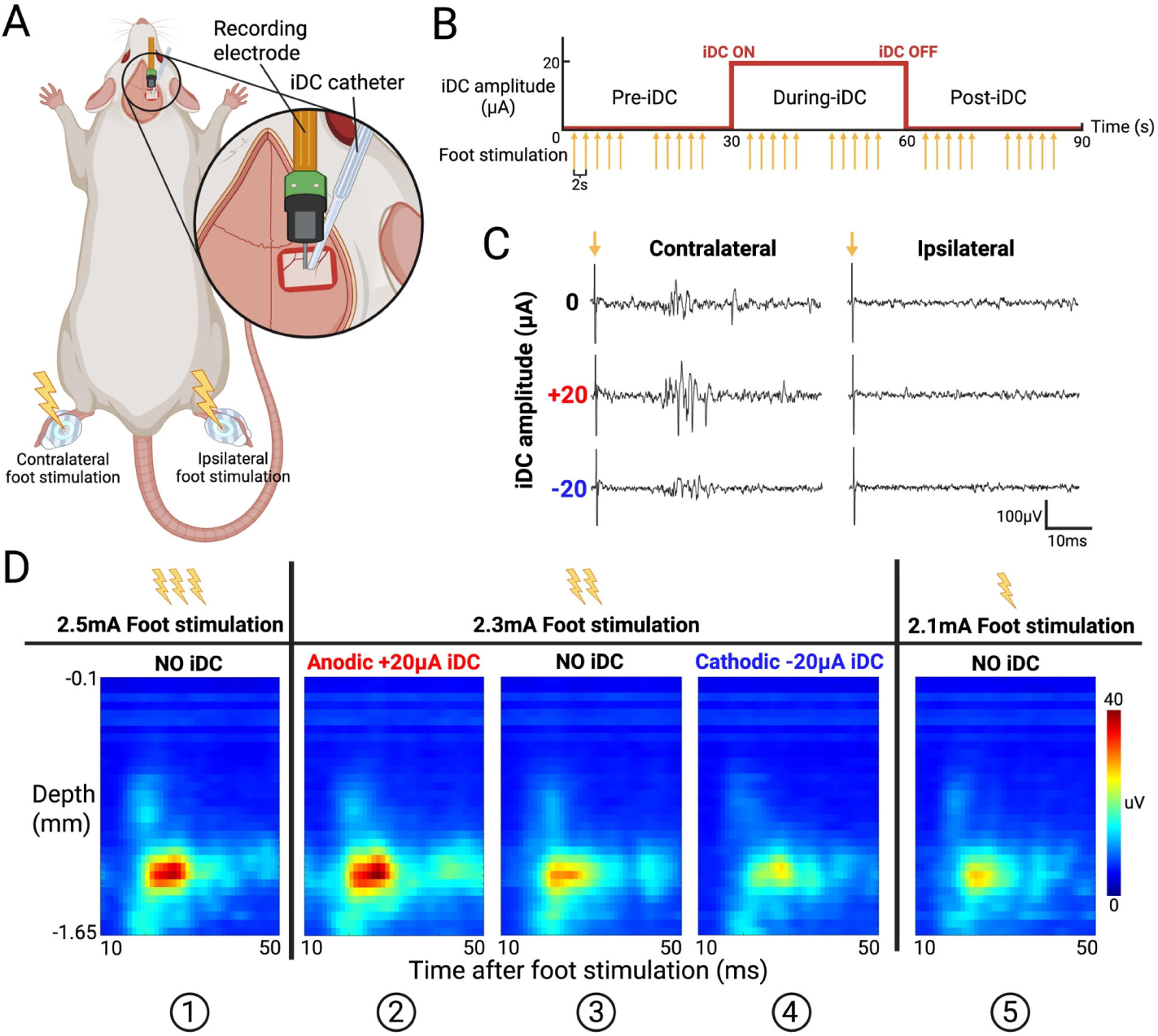
**A**) iDC stimulation catheter, foot stimulation surface electrode, and recording electrode setup used in acute foot stimulation experiment. **B**) iDC (red) and foot shock (yellow) stimulation timeline in each 90s recording session. There are two cycles of 5 repeated foot stimulations per epoch (10 total), each 2s apart. **C**) Example MUA signal recorded from one L5 electrode channel. Evoked responses are mostly generated by the contralateral foot shock, with anodic iDC amplifying the response amplitude and cathodic iDC attenuating it. Arrows indicate the onset of foot shock. **D**) Averaged foot stimulation response heatmaps from one representative experiment. Three center panels (②, ③, ④) show the response to the same foot shock, but with anodic iDC gain amplification on the left (②) and cathodic gain attenuation on the right (④). Note the similarity of iDC-modulated responses to higher and lower amplitude foot stimulation on the corresponding far left (① vs ②) and far right panels (④ vs ⑤).

Figure 4C shows a set of representative MUA responses recorded from a L5 electrode channel. Stimulation of the contralateral foot without iDC produced clear responses that emerged at approximately 20 ms post-stimulus, consistent with the delay associated with somatosensory signal propagation (*36*). When a +20 µA anodic iDC was applied, the evoked responses were noticeably amplified, whereas –20µA cathodic iDC attenuated the response amplitude. As would be expected, ipsilateral stimulation elicited minimal activity with or without iDC stimulation. These observations indicate that iDC modulates the input/output gain of evoked responses in a polarity-dependent manner.

To examine how iDC shifts the gain of evoked responses across cortical depth, we generated time-resolved ESA heat maps from the foot-stimulation MUA (averaged over 10 repetitions per epoch). In these heat maps, the x-axis represents time (10–50 ms window post-stimulus), and the y-axis represents electrode channels arranged by cortical depth (Fig.4D). In this representative experiment, a range of foot stimulation intensities from 2.1 to 2.5 mA was empirically determined to be the effective range intensities eliciting distinct levels of evoked responses. Notably, the heat map for a 2.3 mA foot shock with anodic +20 µA iDC (②) closely resembled that for a 2.5 mA foot shock without iDC (①), while the heat map for 2.3 mA foot shock with cathodic –20 µA iDC (④) was similar to that for 2.1 mA foot shock without iDC (⑤). These comparisons suggest that iDC can shift the effective input/output gain of the local network, with anodic iDC amplifying the evoked response so that a lower-intensity stimulus resembles a higher-intensity one, and cathodic iDC attenuating the response toward that of a lower-intensity stimulus.

To test this gain-shift hypothesis across a broader range of intensities, we constructed evoked-response heat maps for foot-shock amplitudes spanning 0.8–2.8 mA in 0.2 mA steps (Fig. 5A). For each stimulus intensity there were two recordings, one with +20 µA iDC and one with −20 µA iDC, with the NO iDC condition present in the pre-iDC and post-iDC epochs of each recording (Fig. 4B). Thus, each amplitude contributed four NO iDC epochs (two from odd-numbered recordings and two from even-numbered recordings). Because the stimulation parameters and recording configuration were identical within an intensity, we expected the pre-iDC (and post-iDC) responses to be highly similar across cycles. To verify this baseline stability, we first compared pre-iDC heat maps from odd and even cycles by computing a mean squared error (MSE) matrix between even-numbered maps (columns) and odd-numbered maps (rows). As expected, the lowest MSE values clustered along the diagonal (Fig. 5B), indicating that baseline responses were reproducible across repeated recordings at each intensity.

**Figure 5.**
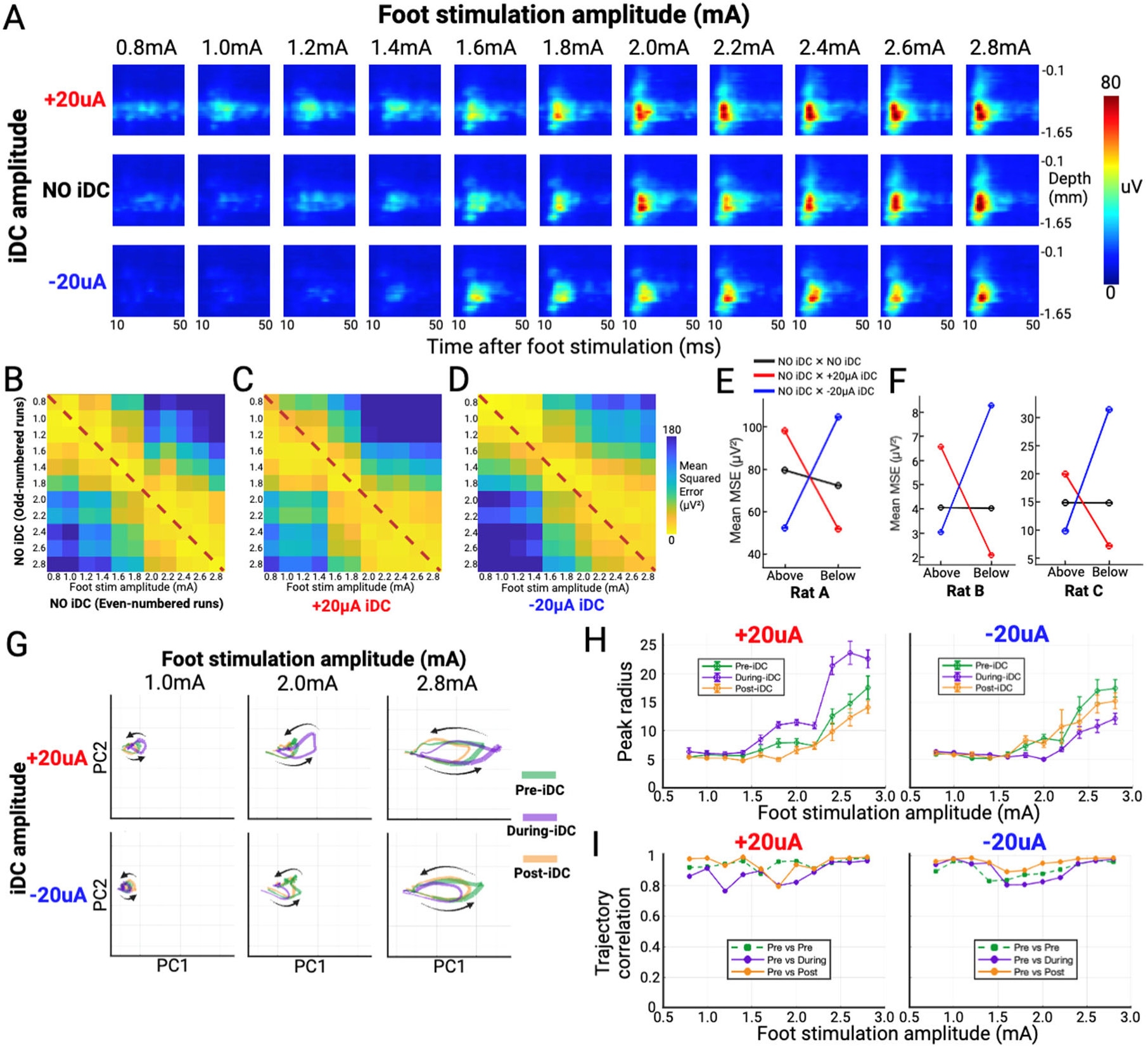
**A**) Depth–time ESA heat maps from a representative experiment delivering a broad range of foot-shock amplitudes (0.8–2.8 mA, 0.2 mA steps) under three iDC conditions: NO iDC (pre-iDC baseline), +20 µA, or −20 µA. **B–D)** Mean squared error (MSE) matrices quantifying similarity between ESA heat maps. Warmer colors indicate lower MSE (greater similarity) while the dashed diagonal marks comparisons at matched stimulus amplitudes. **B)** Even-numbered (x-axis) vs odd-numbered (y-axis) pre-iDC baseline. **C)** +20 µA during-iDC (x-axis) vs pre-iDC baseline (y-axis). **D)** –20 µA during-iDC (x-axis) vs pre-iDC baseline (y-axis). **E)** Mean MSE value above and below the diagonal for matrices **5B, 5C, and 5D**, showing that +20 µA iDC shifts responses toward those evoked by higher-amplitude baseline foot-shocks, whereas −20 µA iDC shifts them toward lower-amplitude shocks. F) Same summary for two additional rats. **G**) Population manifold trajectories (PC1–PC2) of ESA responses for three foot-shock amplitudes for one animal. The common latent space was obtained on all pre-iDC trials across amplitudes. Pre-iDC (green), during-iDC (purple), and post-iDC (orange) mean trajectories are projections into this space. Arrows indicate trajectory direction, and line thickness encodes the normalized across-trial standard deviation of trajectory radius. **H)** Peak trajectory radius (mean ± SEM) in PC1–PC2 as a function of stimulus amplitude, demonstrating bidirectional gain modulation by +20 µA and −20 µA iDC. **I**) Trajectory shape similarity in PC1–PC2 space. For each amplitude, Pearson correlation is computed between pre– and during-iDC mean trajectories (purple), pre– and post-iDC trajectories (orange), and the two baseline pre-iDC blocks (green, pre–pre control), indicating that iDC primarily rescales rather than reshapes the response temporal manifold.

We then used the same MSE framework to evaluate how iDC shifts evoked responses relative to these baselines. Odd-numbered pre-iDC heat maps were compared with +20 µA during-iDC heat maps (Fig. 5C) and with −20 µA during-iDC heat maps (Fig. 5D). Under the +20 µA condition, the lowest MSE values shifted below the diagonal toward the lower left, indicating that responses recorded during anodic iDC more closely resembled baseline heat maps obtained at higher foot-shock intensities. In contrast, under −20 µA conditions, the lowest MSE values shifted above the diagonal and to the right, corresponding to baseline responses evoked by lower stimulation intensities. To summarize these polarity-dependent shifts, we computed the mean MSE above and below the diagonal for all three matrices within each animal. For the representative animal, these summary values showed a clear bias toward lower-intensity baselines for −20 µA iDC and toward higher-intensity baselines for +20 µA iDC (Fig. 5E), and this qualitative pattern was preserved across animals despite inter-animal variability in absolute ESA magnitude (Fig. 5F). Together, the depth-time heat maps and MSE analysis support the interpretation that iDC systematically shifts the effective input/output gain of the local S1HL network.

To examine whether this gain modulation alters the underlying population dynamics or primarily rescales them, we next projected the depth-resolved evoked ESA into a low-dimensional latent space. For each animal, we treated the 10–50 ms post-stimulus ESA at each time bin as a 32-dimensional population vector (one dimension per channel), pooled all pre-iDC trials across foot-shock amplitudes and polarities, and applied PCA to build an animal-specific “column manifold.” In the representative animal shown in Figure 5G**–I**, the first two principal components captured 77.2% of the variance in evoked ESA (PC1 = 69.2%, PC2 = 8.0%), indicating that the dominant stimulus-locked dynamics are well described in the low-dimensional PC1–PC2 plane. When projected into this space, evoked responses at each foot-shock amplitude formed a compact, loop-like trajectory over 10–50 ms post-stimulus. Across amplitudes, +20 µA iDC and −20 µA iDC conditions remained closely aligned with their corresponding pre-iDC trajectories in shape, but with visibly enlarged or shrunken excursions, respectively (Fig. 5G).

To quantify this gain modulation in manifold space, we computed at each time point the radius of the trajectory in PC1–PC2 as the Euclidean norm of the population state, then extracted the peak radius within the 10–50 ms window for each trial. These peak radii were averaged across the 10 trials in each condition (pre-, during-, and post-iDC) and plotted as a function of foot-shock amplitude for +20 µA and −20 µA iDC (Fig. 5H). In the representative animal, for stimulus amplitudes at or above 1.6 mA, +20 µA iDC increased the peak radius relative to baseline with a mean gain ratio of 1.46 ± 0.15 (mean ± SD across 7 amplitudes), whereas −20 µA iDC reduced the peak radius with a mean ratio of 0.73 ± 0.12. After the current was turned off, peak radii returned toward baseline magnitude over the same amplitude range. These manifold-level measurements corroborate the MSE analysis, showing that iDC acts as a polarity-dependent, approximately reversible gain control on the amplitude of the evoked population excursion.

Finally, we asked whether this gain modulation preserved the underlying temporal and geometric “motif” of the evoked trajectories. For each amplitude and condition, we concatenated the mean PC1 and PC2 time courses into a single vector and computed Pearson’s correlation between pre-iDC and during-iDC trajectories, and between pre-iDC and post-iDC trajectories. As a baseline control, we also split the pre-iDC trials into two non-overlapping halves and computed pre–pre correlations at each amplitude. Across amplitudes in the representative animal, pre–during correlations remained high (median r ≈ 0.91; range 0.77-0.98) and were comparable to or slightly below the pre–pre baseline (median r ≈ 0.94; range 0.83-0.98), while pre–post correlations were consistently highest (median r ≈ 0.97; range 0.80-0.99) (Fig. 5I). Thus, even when +20 µA iDC substantially increased manifold radius and −20 µA iDC reduced it, the polarity-dependent changes acted primarily as a *multiplicative gain* on a stable, stimulus-specific response trajectory, rather than deforming them beyond baseline variability.

Together with the MSE analysis of depth–time ESA heat maps, these manifold results indicate that iDC primarily rescales amplitude-specific response trajectories in latent space while preserving their stimulus-specific temporal and spatial motif. In combination with the AIS-centered field model, this supports a picture in which modest, polarity-dependent shifts in deep-layer pyramidal AIS membrane potential act as an effective gain knob on stimulus-locked population dynamics, rather than fundamentally reconfiguring their underlying temporal structure.

### Behavioral Modulation of Tactile Sensitivity Using Von Frey Tests

Having shown in anesthetized cortex that iDC produces polarity-dependent gain changes in spontaneous and stimulus-evoked activity within S1HL, we next asked whether this columnar gain modulation translates into a measurable change in tactile perception in awake rats. The von Frey threshold test provides a quantitative readout of tactile sensitivity during iDC without anesthesia confounds. Because iDC preserves native timing and biases gain rather than directly driving spikes, we predicted a shift in detection thresholds without overt movements in response to iDC alone.

Figure 6A shows the separated-interface headcap system that delivers iDC at the pia while confining faradaic reactions to remote electrolytic reservoirs. Figure 6B shows the block timeline: a constant iDC amplitude applied for 100 s, five von Frey trials at ∼8 s intervals during iDC, then at least 60 s of rest. Amplitudes and paw order were randomized within day. Per block, the 50% withdrawal threshold was estimated with the Simplified Up–Down (SUDO) method (*37*, *38*), then entered into a linear mixed-effects model (see Methods).

**Figure 6.**
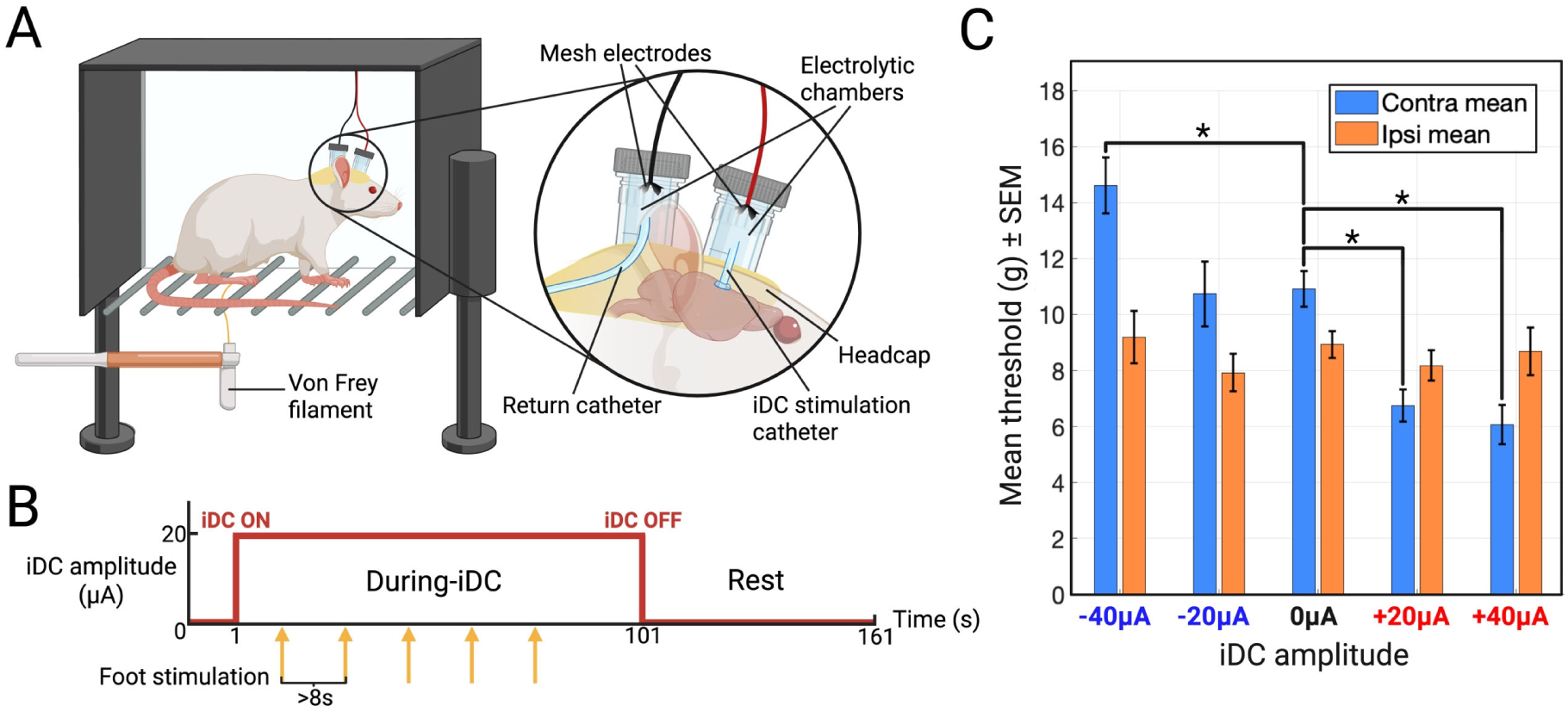
**A**) Chronic headcap for iDC delivery used in von Frey experiments. An agar-gelled aCSF iDC catheter sits on the right S1HL and a return catheter is tunneled to the posterior neck. Both connect to sealed reservoir chambers with stainless-mesh electrodes and a tether to the current source (Keithley 6221). This isolates faradaic reactions to the reservoirs and protects the cortex under the catheters. **B)** Timeline for each von Frey session. A constant iDC amplitude was applied for 100 s per block while five von Frey trials were delivered at ∼8 s intervals, followed by at least 60 s of rest. Amplitudes were randomized within day (−40, −20, 0, +20, +40 µA). Both hind paws were tested with paw order randomized. **C)** von Frey withdrawal thresholds (grams) as mean ± SEM across two rats, blocks and days for each amplitude and paw. Thresholds were computed with the SUDO method from five trials per block. Contralateral to the implant, +20 and +40 µA reduced thresholds and −40 µA increased thresholds. Ipsilateral thresholds showed no significant change. Asterisks mark planned contrasts versus 0 µA within the same paw after Benjamini–Hochberg FDR correction (q < 0.05).

Across both rats, we observed no overt motor response to iDC onset or offset for either anodic or cathodic currents, whereas the withdrawal thresholds to von Frey paw stimulation showed clear polarity-dependent shifts consistent with cortical gain modulation. Data in Fig. 6C are pooled across both animals (mean ± SEM), and statistics reflect a linear mixed-effects model with random intercepts for Rat and Rat:Session. On the contralateral paw, anodic iDC significantly lowered thresholds and cathodic iDC raised them relative to 0 µA. In the mixed-effects model with threshold (g) as the response and amplitude as a categorical factor, +20 µA reduced thresholds by 3.79 g (β = −3.794 g, SE = 0.940, DF = 109, q = 1.35 × 10⁻⁴, 95% CI [−5.657, −1.931], N = 21 blocks) and +40 µA by 4.33 g (β = −4.329 g, SE = 0.971, DF = 109, q = 6.37 × 10⁻⁵, 95% CI [−6.252, −2.405], N = 20). Cathodic −40 µA increased thresholds by 4.21 g (β = +4.213 g, SE = 0.971, DF = 109, q = 6.37 × 10⁻⁵, 95% CI [+2.290, +6.137], N = 20), whereas −20 µA was not significantly different from 0 µA (β = +0.250 g, q = 0.794, 95% CI [−1.644, +2.145], N = 20) (Fig. 6C). This bidirectional, amplitude-dependent pattern mirrors the laminar scaling seen under urethane and is consistent with a gain-control mechanism in the S1HL column that lowers or raises the external force needed to reach a withdrawal criterion.

The ipsilateral paw served as a within-animal control for nonspecific effects such as arousal, tactile habituation, and tethering. Thresholds on the ipsilateral side were unchanged across amplitudes. None of the contrasts versus 0 µA reached significance after Benjamini–Hochberg FDR correction (−40 µA: β = +0.686 g, q = 0.855; −20 µA: β = −0.402 g, q = 0.855; +20 µA: β = −0.144 g, q = 0.855; +40 µA: β = +0.169 g, q = 0.855; DF = 106; N = 19–20 blocks) (Fig. 6C). Both animals performed similarly and showed no overt motor reaction to iDC onset or offset during testing.

Together, these results link cortical modulation to behavior in a somatotopically specific manner. Focal iDC over S1HL decreases contralateral detection thresholds with anodic polarity and increases them with cathodic polarity while the ipsilateral paw remains stable. The polarity and laterality specificity argue that iDC shifts population gain within the targeted cortical columns rather than producing nonspecific arousal or motor effects, providing a causal bridge from the column-scale gain modulation seen in the laminar ESA and manifold analyses to a measurable change in tactile detection behavior.

## DISCUSSION

We selected S1HL as a validation arena because S1 has a well-supported causal role in contralateral tactile perception, and manipulations of S1 can bidirectionally modulate mechanical sensitivity in rodents (*39–41*). Using iDC as a localized, graded perturbation, our electrophysiological recordings and behavioral von Frey tests recapitulated the expected laterality, with contralateral effects and no ipsilateral change, thereby validating iDC as a tool for probing causal relationships in local cortical circuits. Our findings indicate that iDC delivered via a cortical surface microcatheter can bidirectionally bias cortical responses in a controlled and graded manner, modulating local network excitability. Anodic iDC (positive current at the pia) amplified both spontaneous oscillations and sensory-evoked responses, whereas cathodic iDC had the opposite, suppressive effect.

Analysis of neural responses across multiple foot shock amplitudes with and without iDC suggests that iDC can modulate evoked activity in a way that mimics the layer-dependent spatiotemporal response pattern elicited by higher or lower amplitude exogenous input. Consistent with this, PCA-based manifold analyses showed that iDC primarily rescales the excursion of stimulus-specific trajectories in latent space while preserving their temporal and spatial motifs. The ability to modulate “volume” of activity without disturbing the underlying pattern of activity is important to maintaining natural population code used by the brain to communicate between cortical network (*10–12*, *42*, *43*). In this context, the implication is that the saliency of neural response encoded by the overall number of spikes in a population is modulated by iDC, but the underlying structure that represents the information content of the signal remains undisturbed.

### Mechanistic Insights of iDC and Cortical Gain Modulation

The concordance between model predictions and experimental results indicates that field-induced modulation predominantly affects membrane potential near the axon initial segment (AIS), where action potentials are initiated. It suggests that whether that effect was suppressive or excitatory depended on where the cell body was positioned along the layers relative to the dendritic arbor. The vertical length of the dendritic arbor and its position within the electric field determined the membrane potential changes along its length (*2*, *16*, *44*, *45*) and at the soma. Because iDC is quasi-static on neuronal time scales, the induced polarization acts as a steady offset to the membrane potential rather than a rapidly varying drive, biasing the AIS threshold over many integration windows without enforcing phase-locked spiking (*16*, *18*, *46*). The agreement between model and data is consistent with AIS-proximal biasing of dendritic integration, and with the fact that the extracellular electrophysiological recording is dominated by action potentials generated near the AIS/near-soma region, which our laminar ESA readouts capture(*47*, *48*).

We propose that the mechanism by which exogenous electric fields modulate network sensitivity can be understood by recognizing that the axon initial segment (AIS), located near the soma, serves as the integration site for excitatory and inhibitory postsynaptic potentials (EPSPs and IPSPs) originating from pyramidal dendritic synapses (*49*, *50*). We can think of the change in the pyramidal neuron’s AIS membrane potential (Δ*V*_AIS_) as being the sum of the synaptic EPSPs (∑ *EPSP*) minus IPSPs (∑ *IPSP*) and the change in membrane potential imposed by the electric field (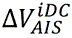):

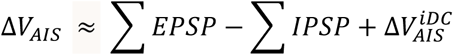

A slight depolarization of the AIS membrane potential (i.e. positive 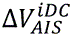) biases the neuron toward excitation, lowering the threshold for excitatory postsynaptic potentials (EPSPs) to trigger an action potential. Conversely, slight hyperpolarization (negative 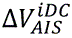) shifts the balance toward inhibition, raising the threshold and making it more difficult for EPSPs to elicit a spike.

### Layer– and Cell-Type Specific Effects

The effects of iDC were not uniform across cortical layers or neuronal cell types. Our in-vivo and previous in-vitro studies (*16*, *18*) both found that under anodic iDC, L5 pyramidal neurons displayed significantly enhanced firing and larger sensory-evoked depolarizations, whereas superficial neurons in L2/3 and putative inhibitory interneurons exhibited comparatively modest responses.

The computational model offers a plausible explanation for this layer-specific modulation, demonstrating that somatodendritic orientation and the differential exposure of neuronal morphologies to the applied electric field govern the extent of voltage gradient “picked up” by neurons (*16*, *18*, *51*, *52*). L5 pyramidal neurons are large projection neurons with long apical dendrites extending toward the cortical surface and somatic compartments positioned near the base of the dendritic volume, with their somatodendritic axis nearly parallel to the radial field emanating from the pia-mounted iDC source. Consequently, these neurons experience a significant voltage drop from dendrite to soma in the presence of iDC, resulting in a higher net membrane polarization at AIS. Additionally, previous studies (*53–56*) indicate that L5 neurons have the highest basal firing rates under urethane anesthesia, whereas L2-4 neurons exhibit sparser spontaneous activity, explaining why iDC modulation produces proportionally larger changes in L5 neuronal activity.

By contrast, smaller cells or those with more symmetric arborizations (such as some interneurons or small excitatory cells in upper layers) likely experience a more uniform electric potential across their structure, and their AIS positions lie near the biphasic zero-crossing of the field, resulting in minimal net polarizations and thus modest modulation by iDC. These findings align with previous work highlighting neuronal morphology and orientation as critical determinants of responsiveness to electric fields (*16*, *18*, *51*, *52*, *57–60*).

Functionally, preferential modulation of L5 pyramidal neurons significantly impacts cortical circuit dynamics. L5 pyramidal neurons constitute major cortical outputs, projecting subcortically and providing excitatory drive to local microcircuits through translaminar synapses. From this perspective iDC neuromodulation can be thought of as “volume control” for the output of a local cortical circuit, consistent with the amplitude-specific ESA scaling and manifold-radius changes we observe in S1HL.

### Comparison with Other Neuromodulation Techniques

The iDC approach provides neuromodulation distinct from, and complementary to, optogenetic and chemogenetics techniques. Optogenetics affords millisecond precision and cell-type specificity but necessitates genetic manipulation and implanted hardware, posing clinical translation challenges (*7*, *8*, *61*). Chemogenetics (e.g., DREADD-based approaches) targets genetically defined neurons through engineered receptors activated by systemic ligands, achieving sustained cell-specific modulation (*62*, *63*). However, chemogenetics modulation incurs inherent delays (minutes) and relies on exogenous ligands, limiting real-time applications (*61*).

Transcranial direct current stimulation (tDCS) produces diffuse sub-millivolt membrane polarization due to the smoothing effect of the skull and scalp, functioning as a broad neuromodulatory influence rather than millimeter-scale focal stimulation(*16*, *51*, *64*). In contrast, iDC provides superior spatial precision and efficacy due to direct placement on the pia mater rather than through the scalp, restricting current spread to targeted cortical columns. Neurons located approximately 1 mm from the iDC catheter displayed negligible modulation, highlighting iDC’s millimeter-scale focality. While this paper investigated the temporal and spatial precision of the iDC modulation, in principle it could be expanded to spatially arbitrary stimulation patterns with broader surface area electrodes.

Lastly, intracortical microstimulation (ICMS) with microelectrodes is a well-established technique for activating neural tissue in a localized manner (e.g., in brain–machine interfaces) (*65*). ICMS typically uses charge-balanced high-frequency pulses to evoke APs phase-locked to pulse presentations, but offers limited ability to produce sustained subthreshold changes or direct inhibition of neural activity (*1*).

By comparison, iDC occupies a middle ground: it uniquely combines spatial specificity and continuous modulation without requiring genetic tools, providing rapid, steady, and reversible neuromodulation suitable for causally probing cortical node functions in behavior and cognition, effectively allowing neuroscientists to “turn up or down the volume” of a cortical node and observe the consequences for circuit dynamics and behavior. Practically, iDC integrates seamlessly with electrophysiological recordings, avoiding electromagnetic interference associated with high-frequency stimulation (*60*, *65*). This compatibility facilitates continuous monitoring of the network’s real-time response to the modulation.

### Future Potential of iDC and Freeform Stimulator Technology

Ionic direct current (iDC) offers precise, bidirectional neuromodulation without the electrochemical damage typical of conventional electrodes. Traditional metal stimulators rely on charge-balanced pulses to avoid tissue degradation but inherently favor excitation and limit sustained inhibition (*19*). In contrast, the electrolyte-filled microcatheter, adapted from the Separated Interface Nerve Electrode (SINE) (*21*), safely delivers true DC by isolating metal components outside the brain. The Freeform Stimulator (FS) implant currently under development further advances this approach by using microfluidic rectification to generate arbitrary ionic waveforms, including direct current, without accumulating charge or contaminants, enabling repeated, long-duration use (*1*, *2*). It is being developed for multichannel and wireless control, supporting complex current steering and spatial field shaping across cortical areas. FS provides a novel tool by enabling chronic application of iDC for long duration behavioral and clinical studies. Clinically, iDC’s gradual, stable modulation is promising for conditions involving abnormal excitability, including epilepsy, stroke, chronic pain, autism, and depression. Together, iDC and FS represent a flexible platform for both research and therapy, enabling high-precision, low-side-effect neuromodulation in both acute and chronic settings.

## MATERIALS AND METHODS

### Acute Experimental Procedures

#### Animals and Surgical Preparation

All animal procedures were approved by the Institutional Animal Care and Use Committee (IACUC) of Johns Hopkins University and conducted in compliance with NIH guidelines. Adult Sprague–Dawley rats (n=7; 4 males, 3 females; 450–600 g) were housed with a 12 h light/dark cycle and given food and water ad libitum. Animals were anesthetized with urethane (1.4 g/kg, intraperitoneal injection). Anesthetic depth was monitored regularly during surgery via heart rate, foot-pinch reflex, and blink response. Body temperature was maintained at 37 ± 0.5 °C using a feedback-controlled heating pad.

A craniotomy was performed to expose the hindlimb region of the primary somatosensory cortex (S1HL) in the right hemisphere (coordinates: posterior bregma −0.5 to −1.5 mm, lateral 2.0 to 3.5 mm). The dura mater was carefully removed, and a 32-channel single-shank microelectrode array (NeuroNexus A1×32-6mm-50-177-Z32) was vertically inserted ∼1.65 mm into the cortex, spanning all layers with its 1.55 mm contact array (Fig. 1DE). A stainless-steel reference electrode was positioned between the skin and muscle tissue on the contralateral side of the skull. S1HL localization was confirmed by gently brushing the contralateral hindfoot and verifying robust multi-unit activity (MUA) responses.

#### Stimulation and Recording

A 28-gauge non-conducting MicroFil flexible needle (250 µm inner diameter) was filled with agar-gelled artificial cerebrospinal fluid (aCSF). The microcatheter tip was gently placed on top of the pia mater within S1HL, adjacent to the inserted recording probe (∼0.2 mm), and oriented perpendicular to the cortical surface (Fig. 1DE). The microcatheter was connected to an external constant current stimulator (Keithley 6221) through a hypodermic needle on top. This separated-interface microcatheter setup provides a temporary solution to safely deliver iDC currents and isolates the metal interface from brain tissue to temporarily mimic the function of the Freeform Stimulator. A separate hypodermic needle served as the return electrode and was placed subcutaneously at the animal’s tail base. Prior testing using pH-sensitive dyes confirmed that electrochemical byproducts at the metallic portion of the circuit were contained away from the cortex for the duration of the experiments.

Neural signals were amplified (×1000) and digitized at 24.414 kHz using Tucker-Davis Technologies hardware (Subject Interface and processor) and Synapse software. For MUA extraction, recordings were bandpass filtered between 300 and 5000 Hz. Both spontaneous and foot-stimulation-evoked MUA recordings were conducted in approximately 90-second sessions, segmented into 30 s epochs for pre-iDC, during-iDC, and post-iDC intervals (Fig. 1F). iDC amplitudes varied randomly across sessions, ranging from –50 to +50 µA.

For spontaneous activity recordings, to minimize artifacts due to edge effects and stimulation onset/offset, a 500 ms interval at both the start and end of each 30-second epoch was excluded, yielding an effective analysis window of 29 seconds per condition (pre-iDC: 0.5–29.5 s; during-iDC: 30.5–59.5 s; post-iDC: 60.5–89.5 s).

Evoked responses were elicited by delivering cathodic foot stimulation pulses (200 µs, 0.5 to 2.8 mA; Model 2100, A-M Systems, Sequim, WA) every 2 seconds, repeated 10 times per epoch (3 epochs × 10 stimulations per recording; Fig. 4B). Stimuli were applied via 3M Red Dot ECG electrodes (Model 2560), with the cathode on the plantar and the anode on the dorsal surface of the contralateral hindfoot (Fig. 4A). A control stimulus was applied to a different dermatome area (e.g., ipsilateral foot) to confirm the specificity of iDC effects on contralateral S1HL. Evoked MUA responses were evaluated in the 10–50 ms post-stimulus window.

To evaluate the spatial resolution of electric field effects, the iDC microcatheter was positioned at incremental horizontal offsets (0.2, 0.55, 0.90, and 1.25 mm) from the recording electrode across 5 animals (3 males, 2 females; Fig. 3DE), and spontaneous activity was recorded at various iDC amplitudes for each offset.

### Acute Experiment Data Analysis

Having acquired spontaneous and evoked MUA under randomized iDC amplitudes, we next applied custom MATLAB (The MathWorks Inc, R2023b) signal-processing pipelines to quantify gain changes due to iDC stimulation.

#### Entire Spiking Activity (ESA) Analysis

Both spontaneous and foot stimulation evoked MUA signals were processed using the Entire Spiking Activity (ESA) method over their relevant time window (e.g., 29 s for spontaneous activity or 10–50 ms for evoked responses) (*23–25*). MUA signals were full-wave rectified and bidirectionally low-pass filtered with a Gaussian kernel (σ = 1 ms, kernel length = 6σ) (Fig. 2A). The ESA method preserves activity from smaller neurons by avoiding threshold-based spike detection, allowing more robust quantification under varying signal-to-noise conditions (*23*, *24*).

To visualize spontaneous activity across cortical depth, ESA signals from each channel were segmented into 50-ms non-overlapping bins across the 29 s analysis window, then vertically arranged by cortical depth to generate time-resolved heatmaps (Fig. 2B). For foot stimulation evoked activity, the MUA responses were first aligned to each stimulus onset, and a 5-ms post-stimulus “artifact” window was excluded. To ensure stable edge behavior, the retained 10–50 ms post-stimulus effective analysis window was padded by 5 ms on either side, yielding an extended 5–55 ms segment. This segment is then processed by the ESA method, cropped back to the 10–50 ms window, and divided into 1 ms bins. ESA segments from all 32 channels were depth-aligned and averaged over 10 repetitions to create one response heatmap per epoch (Fig. 4D**, 5A**).

To quantify spontaneous activity changes, spontaneous ESA signals from each channel were also averaged over the 29 s during-iDC epoch and normalized to baseline (pre-iDC epoch) to yield a relative percent change value. These values were calculated across 12 iDC amplitudes (–50 to +50 µA) and arranged by channel depth to form the relative percent change table (Fig.2C). Recording channels were also assigned by depth into four cortical layers (2/3, 4, 5, and 6). Within each experimental cycle (one animal per cycle, 12 iDC amplitudes), we averaged the channel-level relative percent changes (during– vs. pre-iDC or post– vs. pre-iDC) for each layer and then plotted the mean ± SEM across 7 animals as line graphs with error bars (Fig. 2DE).

#### PCA-Derived Power Spectral Density Curves

To confirm that iDC does not disrupt intrinsic oscillatory dynamics, we analyzed raw (unfiltered) spontaneous activity from all 32 channels and 12 iDC amplitudes (–50 µA to +50 µA). Data segments (pre-iDC: 10.5–29.5 s; during-iDC: 40.5–59.5 s) were reduced via principal component analysis (PCA), with the first principal component (PC1) serving as a representative “column signal.” Power spectral density (PSD) was calculated using a median short-time Fourier transform (STFT; 2 s Hamming windows, 0.5 s overlap), with spectral power derived from the median squared magnitude (|STFT|²) over time. Spectra were interpolated onto a log-spaced frequency axis (0.5–3 kHz), smoothed with a 10-point moving average, and converted to decibels (dB). Mean ± SEM PSD curves across three animals (1 male, 2 females) were generated for the pre-iDC baseline (black shading and trace) and each iDC amplitude condition (colored shading and traces) (Fig. 2F).

#### Mean Squared Error (MSE) Analysis of Evoked-Response Heatmaps

To quantify changes in the gain of foot-stimulation evoked responses, we computed pairwise mean squared error (MSE) matrices comparing heat maps from odd-versus even-numbered baseline cycles and between baseline and ±20 μA iDC cycles (Fig. 5B**-D**). For each pair of sessions (indexed along the matrix axes), the two response heatmaps were vectorized, and MSE was computed as the average squared difference (in µV²). Diagonal elements of the matrices (self-comparisons) served as controls. To summarize polarity-dependent shifts, we computed the average MSE above the diagonal (indicating similarity to lower-intensity baselines) and below the diagonal (indicating similarity to higher-intensity baselines) for all matrices in each animal (Fig. 5EF).

#### PCA Manifold Analysis of Evoked ESA Responses

For manifold analysis of stimulus-evoked responses, we used the same artifact-handled ESA preprocessing described above to obtain 10–50 ms post-stimulus ESA snippets (1 ms bins) from all 32 channels. For each recording session (one foot-shock amplitude and one iDC polarity), this yielded 30 trials (10 pre-iDC, 10 during-iDC, 10 post-iDC) arranged as a tensor of size *trials × time × channels* (30 × 40 × 32) in units of µV.

For each animal, we constructed a single latent space by pooling all pre-iDC trials across foot-shock amplitudes and iDC polarities. ESA snippets from these pre-iDC trials were reshaped into a matrix of samples × channels, z-scored across samples for each channel, and subjected to principal component analysis (PCA). The resulting loading matrix defined a common “column manifold” for that animal. The same pre-iDC mean and standard deviation were then used to z-score every trial (pre-, during-, and post-iDC) before projection into this PCA space. For visualization and quantification, we retained the first two principal components (PC1–PC2), which together captured the majority of variance in the evoked ESA (Fig. 5G**–I**).

For each foot-shock amplitude and iDC condition (pre-, during-, post-iDC), PC scores were reshaped back to *trials × time × PCs* and averaged over the 10 trials to obtain a mean trajectory in latent space (*time × PCs*). Trajectories in Fig. 5G show PC1 on the x-axis and PC2 on the y-axis, with arrows indicating the direction of increasing post-stimulus time. Line thickness encodes the normalized across-trial standard deviation of trajectory radius (see below), providing a visual readout of trial-to-trial variability along the manifold.

#### Trajectory Magnitude and Shape Metrics in Latent Space

To quantify how iDC scales the evoked manifold excursion, we computed a trajectory “radius” at each time point as the Euclidean norm in the PC1–PC2 plane. For each trial, the peak radius within the 10–50 ms window was extracted and then averaged across the 10 trials in that condition (pre-, during-, or post-iDC) to yield a mean peak radius and its standard error. Peak-radius versus stimulus-amplitude curves (mean ± SEM) were plotted separately for +20 µA and −20 µA iDC to quantify bidirectional gain modulation in manifold space (Fig. 5H).

To assess whether iDC preserved the temporal/geometry “motif” of the evoked manifold, we compared the shapes of mean trajectories across conditions. For each foot-shock amplitude and condition, the mean PC1 and PC2 time courses were concatenated into a single vector, and Pearson’s correlation coefficient was computed between pre-iDC and during-iDC trajectories, and between pre-iDC and post-iDC trajectories at the same amplitude. As a baseline control for trajectory-shape reliability, for each amplitude the 10 pre-iDC trials were also split into two non-overlapping subsets of 5 trials each, separate mean trajectories were computed for the two subsets, and a pre–pre correlation was calculated using the same procedure. Because Pearson’s correlation is invariant to overall scaling, these metrics report trajectory-shape similarity independent of changes in gain. Pre–during, pre–post, and pre–pre trajectory-shape correlations were plotted as a function of stimulus amplitude for +20 µA and −20 µA iDC (Fig. 5I), providing a manifold-level measure of how well iDC preserved the stimulus-specific response motif while modulating its magnitude.

### Computational Modeling

#### iDC Electric Field Spread

The model first establishes a two-dimensional grid representing a 2.2 × 2.2 mm rat S1 cortex cross-section in MATLAB. The extracellular potential (Vₑ) at every point in the cross-section was computed by superposition of point-source contributions from a 0.25 mm diameter circular disk electrode (mimicking the iDC catheter) placed on the pia (Fig. 3A). Each point source at radius r contributes:

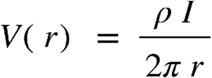

where ρ = 5 × 10^3^ Ωꞏmm is the in vivo measured tissue resistivity (see **Supplemental Fig. 2**) and I is the applied current (i.e., ±20 µA). We model the catheter interface to the pia as an isotropic conductive half-space, so 2πr indicates the hemispherical spread of voltage, with the pia treated as the boundary interface.

#### Neuron-Rod Representation and Mirror Estimate

We represent each cell as a vertical “rod” whose length matches its dendritic span, and assign a discrete soma depth (black markers) to approximate the electrotonic center (Fig. 3B). Eight most common excitatory types (L2 pyramidal, L3 pyramidal, L4 star pyramidal, L4ss: L4 spiny stellate cell, L5 slender-tufted pyramidal, L5 thick-tufted pyramidal, L6 corticocortical pyramidal, L6 corticothalamic pyramidal (*29*, *30*, *32*)) and two inhibitory types (basket and Martinotti (*31*)) were parameterized by empirically derived top and bottom dendritic extents and soma positions. Each rod type was evenly spanned across the cortex cross-section. For each neuron rod, we compute the “mirror estimate,” defined as:

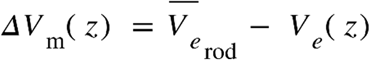

where 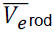 is the mean extracellular potential across the rod’s depth (*2*, *33*, *34*). This metric serves as a DC analog of the extracellular activation function, predicting net transmembrane polarization.

#### Population-Level Heatmap via Weighted Sum Interpolation

To approximate population multiunit activity, rods from each excitatory neuron type were randomly scattered laterally across one cortex cross-section in proportion to their relative densities(*30*). Soma potential changes ΔV_m_ were extracted and placed on a finer 200×200 grid, then summed with Gaussian weighting (σ=0.1 mm) to reflect local neuronal activity:

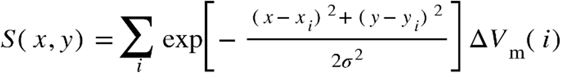

The resulting 2D heatmap displays the spatial distribution of net soma polarization under anodic or cathodic iDC, which serves as an approximation of the activity change that would be detected by the recording electrode used in our in vivo setup (Fig. 3C).

The complete code can be found in https://github.com/RunmingTonyWang/iDC-cortex-model

### Behavioral iDC Experiments in Awake Rats

#### Animals

Adult Sprague–Dawley rats were used (n = 2, one male and one female, 450–550 g). Procedures were approved by the Johns Hopkins Animal Care and Use Committee. Animals were housed on a 12:12 h light–dark cycle with ad libitum food and water.

#### Surgery and Chronic iDC Catheter Implantation

Surgical preparation and craniotomy followed the acute procedures described above. Anesthesia was induced with isoflurane at 2.5% and maintained at 1.5% in oxygen. Depth was monitored by heart rate, pedal withdrawal, and blink reflex. Body temperature was maintained at 37 ± 0.5 °C using a feedback heating pad. A right hemisphere craniotomy exposed S1HL cortex (posterior bregma −0.5 to −1.5 mm, lateral 2.0 to 3.5 mm). The dura was opened as in acute experiments, and the pia was preserved.

A microfluidic iDC catheter was positioned on the pial surface over S1HL. For the chronic build, the outlet iDC catheter was a polyethylene tube (300 um ID) filled with agar-gelled aCSF, oriented perpendicularly to the cortical surface (Fig. 6A). A return catheter of identical construction was tunneled subcutaneously to the posterior neck and oriented away from the head. Catheters were secured with dental acrylic anchored to skull screws. Two custom 3D-printed chambers were affixed at the skull and neck sites and filled with liquid aCSF agar to serve as electrolytic reservoirs (Fig. 6A). Each chamber contained a folded stainless steel mesh electrode with a tethered lead exiting through a sealed cap. This architecture confines faradaic reactions to the remote metal–agar interfaces and prevents electrochemical byproducts from entering the iDC microcatheters. pH-sensitive dye tests confirmed that pH changes at the metal–agar sites did not reach either microcatheters over the session durations used here.

#### Postoperative Care and Habituation

Meloxicam was administered at 1 mg/kg subcutaneously. Animals recovered on a warm pad until fully ambulatory. Postoperative care included twice daily monitoring of the incision, cap integrity, and general health for at least seven days. After recovery, animals underwent at least seven days of daily handling and tether habituation in the testing room.

#### iDC Stimulation During Behavior

During testing, iDC was delivered as a constant amplitude via a current source (Keithley 6221) connected to both steel mesh electrodes through a pair of lightweight cables with slack. Tested amplitudes were −40, −20, 0, +20, and +40 µA. For each behavioral block, iDC was turned on for 100 seconds and all five von Frey trials in that block were performed while iDC was on. At least 60 seconds of rest without stimulation followed each block (Fig. 6B).

#### von Frey Assay and SUDO Thresholding

Mechanical withdrawal thresholds were measured on an elevated mesh platform using monofilaments (Touch Test®; forces 0.008–300 g). The filament was applied perpendicularly to the plantar surface until it just bent and was held for about 1–2 s. Trials were separated by at least 8 s. Thresholds were estimated using the Simplified Up–Down (SUDO) method (*37*, *38*): starting at a pre-specified filament, an observed withdrawal (“X”) resulted in stepping one filament lower for the next trial; no withdrawal (“O”) stepped one filament higher. After the fifth scored trial, the 50% withdrawal threshold index was defined as N5 ± 0.5 (−0.5 for “X”, +0.5 for “O”), and the force was computed as the geometric mean of the two adjacent filaments at that index. Sessions with <5 valid trials or excessive noncompliance were excluded a priori.

#### Experimental Design and Statistical Analysis

The design was within subject and repeated measures. Both hind paws were tested in each rat. Paw order was randomized within day. iDC amplitudes were randomized within paw and day and 0 µA baseline was included each day. Approximately ten blocks were collected per paw per day across about seven experimental days per rat. The experimenter was blinded to iDC amplitude.

Analyses were performed in MATLAB. For each paw, a linear mixed effects model was fit with withdrawal threshold in grams as the response, iDC amplitude as a fixed effect, and random intercepts for Rat and for Session nested within Rat:

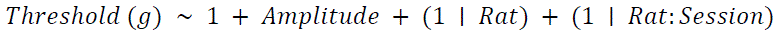

The 0 µA condition was the reference. Planned contrasts compared each amplitude with 0 µA using Wald t-tests with Satterthwaite degrees of freedom. The false discovery rate across amplitudes within each paw was controlled with the Benjamini–Hochberg FDR procedure. Adjusted q less than 0.05 was considered significant. For visualization, Figure 6C shows mean ± SEM thresholds for each amplitude and paw across both rats.

#### Safety and Headcap Maintenance

To keep electrochemical byproducts confined to the reservoirs, the electrolytic gel in both remote chambers was replaced after each experimental day. Catheter patency, chamber seals, and headcap rigidity were verified before testing each day.

## ACKNOWLEDGEMENTS

We thank Grace Foxworthy, W. Mitchel Thomas, Celia Fernandez Brillet, and Katherine Mueller for technical support and helpful discussions. We thank Jiyoon Beris, and Roshin Varghese for assistance with behavioral data collection. Figures were assembled with BioRender (https://BioRender.com/zho7ddg).

## Funding

Funding was provided by the National Institutes of Health (NIH) through the support of grants to G.F. (NIH R01NS110893).

## Author contributions

Conceptualization: R.W. and G.F.

Methodology: R.W. and G.F.

Investigation: R.W. and G.F.

Software: R.W. and G.F.

Formal analysis: R.W. and G.F.

Visualization: R.W. and G.F.

Writing – original draft: R.W. and G.F.

Writing – review & editing: R.W. and G.F.

Supervision: G.F.

Funding acquisition: G.F.

## Competing interests

The authors declare no competing interests.

## Data and materials availability

All data needed to evaluate the conclusions in the paper are present in the paper and/or the Supplementary Materials. Code for computational model is available at: https://github.com/RunmingTonyWang/iDC-cortex-model. The study did not generate new unique reagents. Due to size constraints, all raw electrophysiology recordings and custom analysis code will be provided to editors and reviewers upon request during peer review and deposited with a DOI upon acceptance.

## SUPPLEMENTARY MATERIALS

### Control Experiment for Reference-Wire Artifacts in ESA Metrics

To confirm that reference-wire artifacts did not influence our ESA measurements, we performed control recordings in which anodic and cathodic iDC (±30 µA) were applied using identical stimulation and recording configurations in both living and euthanized rats. ESA relative percent changes during-vs.-before iDC were calculated for both conditions. Anodic and cathodic stimulation robustly altered ESA in layer 5 channels of the alive rat, whereas no change was observed across channels in the euthanized rat (**Supplemental Fig. 1**). These results verify that ESA metrics reported here were unaffected by stimulation artifacts.

**Supplemental Figure 1.**
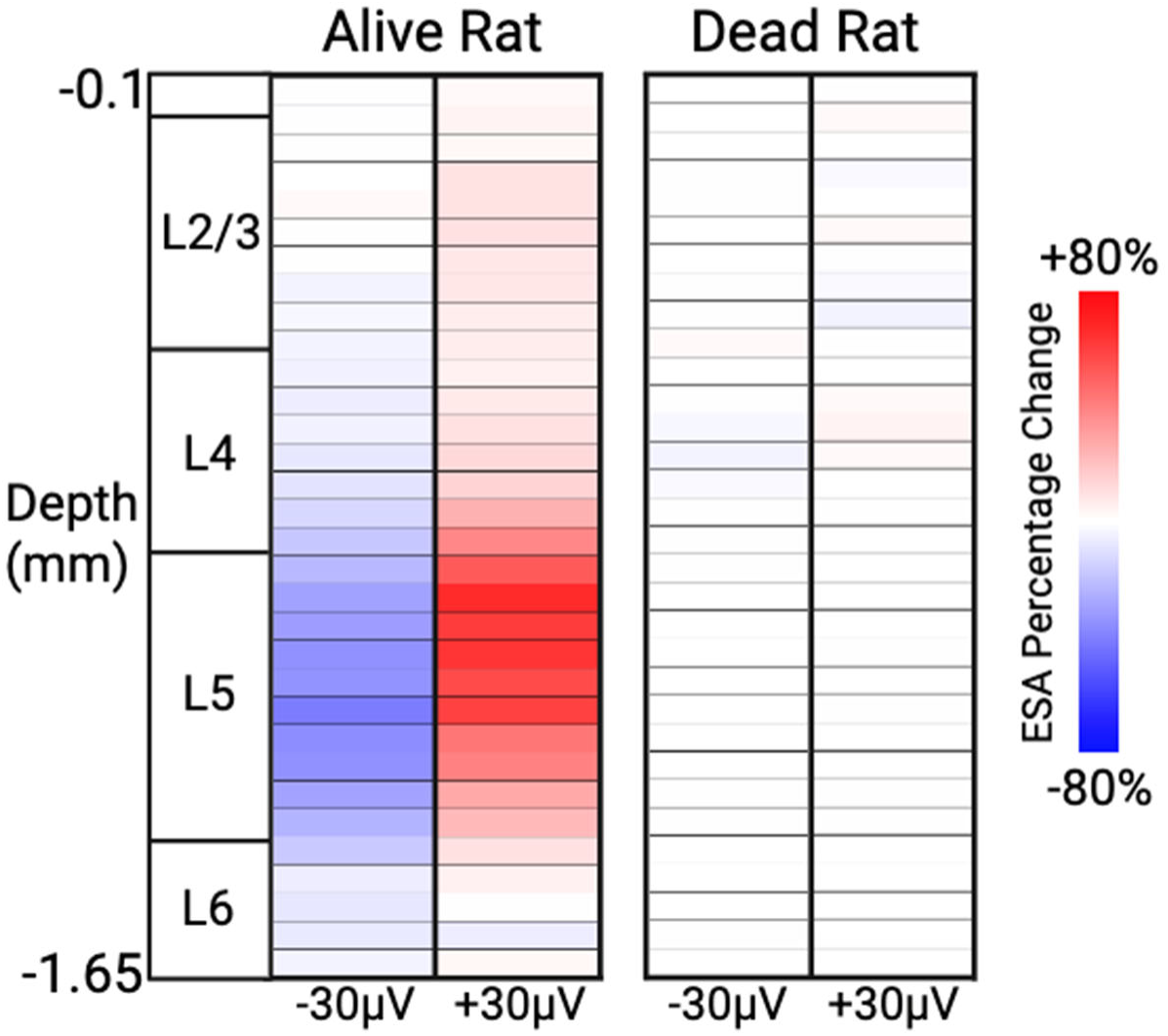
Relative percent change in average ESA across recording channels for alive and euthanized rats.

### Electric Field Mapping In Vivo

To validate the tissue resistivity parameter (ρ) used in our computational model, we delivered brief cathodic pulses (200 µs, –100 µA) through the iDC microcatheter placed at multiple lateral distances (0.25–2.6 mm) from the microelectrode array. Peak voltage amplitudes were recorded from all 32 channels spanning cortical depths (∼0.10–1.65 mm) and lateral offsets (∼0.25–2.55 mm), generating a two-dimensional voltage map of the induced electric field (**Supplemental Fig. 2**). We optimized the resistivity parameter by matching model-predicted voltages to experimentally recorded values, yielding a best-fit resistivity of approximately 5 × 10³ Ωꞏmm, which was subsequently used in our computational model.

**Supplemental Figure 2.**
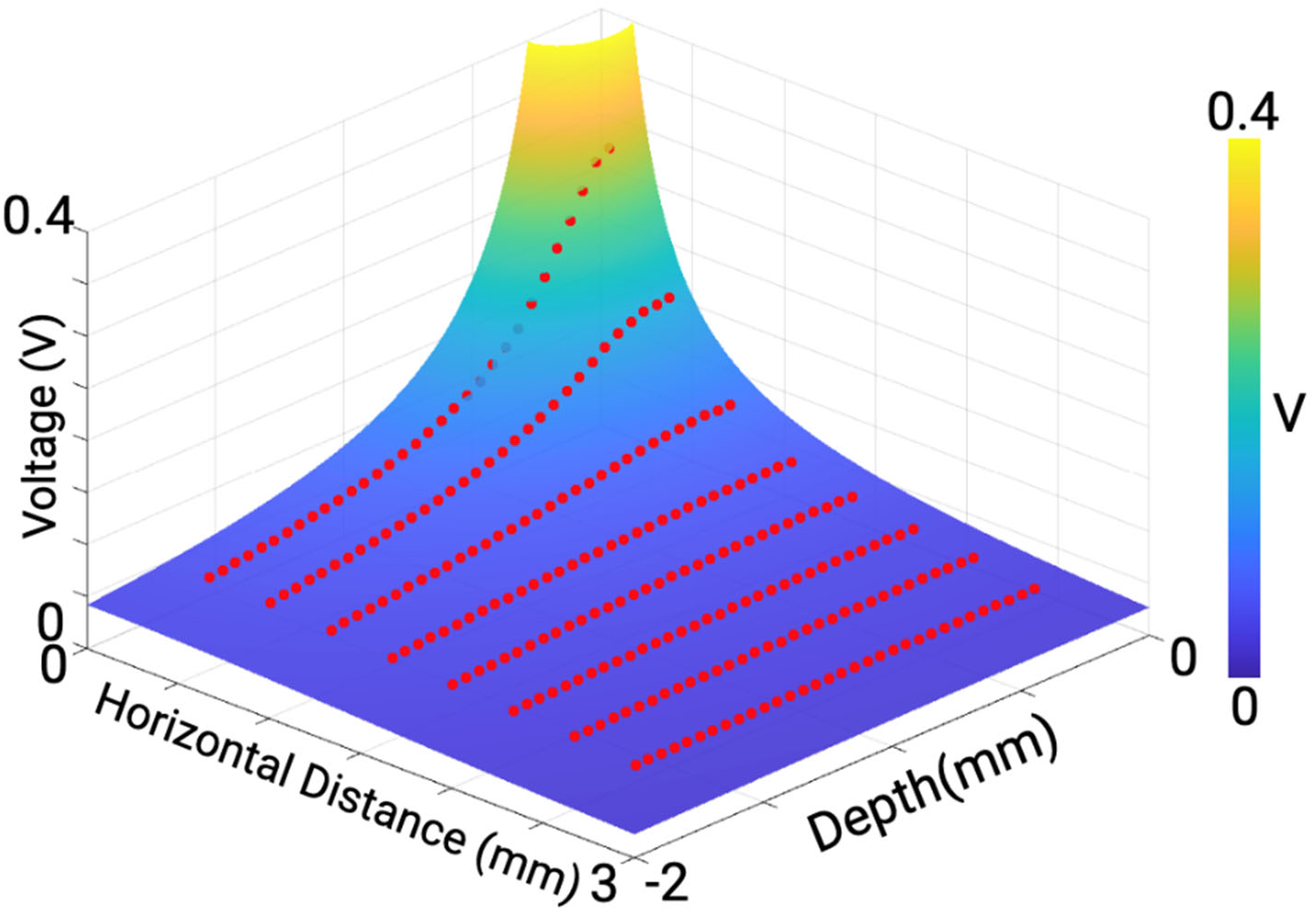
Measured voltages (red dots) overlaid on model-predicted extracellular voltage spread.

## Notes

https://github.com/RunmingTonyWang/iDC-cortex-model

### Competing Interest Statement

The authors have declared no competing interest.

### Summary of Updates

We conducted analysis that strongly suggests that iDC modulates the amplitude of the neural responses (number of evoked spikes) without affecting the population manifold dynamics. (Fig. 5)

